# Oligo-PROTAC strategy for cell-selective and targeted degradation of activated STAT3

**DOI:** 10.1101/2023.08.01.551552

**Authors:** Jeremy Hall, Zhuoran Zhang, Dongfang Wang, Supriyo Bhattacharya, Marice Alcantara, Yong Liang, Piotr Swiderski, Stephen Forman, Larry Kwak, Nagarajan Vaidehi, Marcin Kortylewski

## Abstract

Decoy-oligodeoxynucleotides (D-ODNs) can target undruggable transcription factors, such as STAT3. However, challenges in D-ODN delivery and potency hampered their translation. To overcome these limitations, we conjugated STAT3-specific D-ODN to thalidomide (Tha), a known ligand to cereblon (CRBN, a component of E3 ubiquitin ligase) to generate a proteolysis-targeting chimera (STAT3D^PROTAC^). STAT3D^PROTAC^ downregulated STAT3, but not STAT1 or STAT5, in target cells. Computational modeling of the STAT3D^PROTAC^ ternary complex predicted two surface lysines on STAT3, K601 and K626 as potential ubiquitination sites for the PROTAC bound E3 ligase. Accordingly, K601/K626 point mutations in STAT3, as well as proteasome inhibitors, and CRBN deletion alleviated STAT3D^PROTAC^ effect. Next, we conjugated STAT3D^PROTAC^ to a CpG ligand targeting Toll-like receptor 9 (TLR9) to generate myeloid/B-cell-selective C-STAT3D^PROTAC^ conjugate. Naked C-STAT3D^PROTAC^ was spontaneously internalized by TLR9^+^ myeloid cells, B cells as well as human Ly18 and mouse A20 lymphoma cells, but not by T cells. C-STAT3D^PROTAC^ decreased STAT3 levels to 50% at 250 nM and over 85% at 2 µM dosing in myeloid cells. We also observed significantly improved downregulation of STAT3 target genes involved in lymphoma cell proliferation and/or survival (*BCL2L1, CCND2, MYC*). Finally, we assessed the antitumor efficacy of C-STAT3D^PROTAC^ compared to C-STAT3D or scrambled control (C-SCR) against human lymphoma xenotransplants. Local C-STAT3D^PROTAC^ administration triggered lymphoma regression while control treatments had limited effects. Our results underscore feasibility of using PROTAC strategy for cell-selective, decoy oligonucleotide-based targeting of STAT3 and potentially other tumorigenic transcription factors for cancer therapy.

## INTRODUCTION

Signal transducer and activator of transcription 3 (STAT3) is a transcription factor (TF) and prominent oncogene responsible for tumorigenesis and immune evasion associated with poor prognosis in a variety of human cancers.^1–3^ STAT3 is activated by a variety of upstream cytokine and growth factor receptor-associated tyrosine kinases such as Janus kinase (JAK1, JAK2) or oncogenic Src and Abl kinases, resulting in the formation of homo-or hetero-dimers which are then translocated to the nucleus to initiate downstream gene expression.^1,4^ Abnormal STAT3 signaling in tumor and tumor-associated myeloid cells has been shown to affect the regulation of genes relevant to such cellular functions as angiogenesis, cell signaling, immunosuppression, inflammation, proliferation, and metastasis. As a result, STAT3 has emerged as a distinctly unique and attractive target in cancer therapy.^2,5,6^ However, due to the lack of kinase domain and largely planar surface area for protein-protein interactions, TFs like STAT3 are challenging pharmacologic targets.^5,7,8^ Synthetic oligonucleotides such as STAT3 decoy DNA or antisense oligonucleotides (ASOs) have shown promise in clinical trials and were well tolerated in patients.^9–11^ However, except for hepatocyte targeting, lack of cell-selective delivery strategies remains a key challenge for the majority of oligonucleotide therapeutics.^12^ Broad and non-cell selective STAT3 inhibition is likely to result in conflicting effects on the immune cell network, thereby limiting the long-term antitumor immune responses. This is partly due to STAT3 role in the expansion of cytotoxic CD8 T cells in cancer patients^13^ and in the development and maintenance of memory T cells^14^. To overcome these limitations, we previously developed a strategy to deliver oligonucleotide-based STAT3 inhibitors, such as siRNA, ASO or decoy oligodeoxynucleotides, specifically into tumor-associated myeloid cells, B cells and some cancer cells.^15^ Conjugation of STAT3 decoy to toll-like receptor 9 (TLR9) ligands, CpG oligonucleotides, facilitated targeting of TLR9^+^ immune and cancer cells, prompting immune activation and antitumor responses.^15^ CpG-STAT3 decoy conjugate (CpG-STAT3D) was effective in delivering decoy molecules into human and mouse dendritic cells (DCs), macrophages, and myeloid-derived suppressor cells (MDSCs) and also into myeloid leukemia or B cell lymphoma cells in mice.^16,17^ More recently, we successfully adopted CpG-Decoy strategy for targeting canonical and non-canonical NF-κB signaling specifically in human and mouse B cell-lymphoma cells in vivo.^18^ The antitumor efficacy of both STAT3- and NF-kB-specific CpG-Decoy strategies resulted mainly from the induction of myeloid or B cell differentiation driving the activation of antitumor immune responses.^17,18^ However, the direct cytotoxic effects of decoy molecules against rapidly proliferating cancer cells may have been limited by the reversibility of dose-dependent target inhibition.

Over the last 20 years, proteolysis targeting chimeras (PROTACs) have emerged as a unique modality to target and degrade an intracellular protein of interest (POI) by utilizing E3 ubiquitin ligases for proteasomal degradation.^19,20^ PROTAC consist typically of two small molecules, one which recruits the E3 ligase and another that binds to the POI, connected by a linker molecule.^21^ The chemical nature and length of the linker are critical factors in defining an effective interaction between the POI and the E3 complex.^22^ Since PROTACs do not require high affinity binding to the target, they have potential to extend their inhibitory effect to yet “undruggable” TFs and also yield target specificity.^20^ In addition, the stability of PROTACs enables recycling of the same molecule to degrade multiple copies of the target protein in a catalytical process, maximizing their potency.^23^ PROTAC designs commonly utilize thalidomide or lenalidomide, related immunomodulatory drugs which can recruit cereblon (CRBN) protein within the E3 ligase complex.^21,24^ Such small molecule PROTACs demonstrated ability to degrade multiple oncoproteins, including AKT, BRD4 or EGFR but only lately were tested for STAT3 targeting.^20,24,25^ Recently developed small molecule SD-36 is a lenalidomide-based STAT3 degrader. SD-36 showed activity against acute myeloid leukemia (AML) and large-cell lymphoma cells in vitro and in immunodeficient mice but lacks cell-selectivity critical for generating effective antitumor immune responses.^25,26^

Here, we outline a rational STAT3 Oligo-PROTAC design based on a structural modeling of interactions between a decoy-bound protein target and E3 ligase complex. Coupled with TLR9-directed delivery strategy, our approach allows for cell-targeted and STAT3-selective degradation to improve antitumor efficacy and safety.

## RESULTS

### Oligo-PROTAC design for targeted degradation of STAT3

In order to generate a STAT3 proteolysis-targeting chimeric ODN, we equipped CpG-STAT3 Decoy with thalidomide molecule attached using a propandiol linker to the 3’ end of the double-stranded decoy hairpin (Figure 1A and Supplemental Figure S1).^16,17,27^ Thalidomide, as a ligand for cereblon (CRBN) protein, could facilitate interaction between an E3 ligase complex and the STAT3 protein dimer bound to the decoy conjugate. This hypothesis was supported by in silico analysis using a combination of molecular modeling and molecular dynamics (MD) simulations in solution state. As shown in Figure 1B, the 3’ end-located thalidomide can bind to the tri-Trp pocket of CRBN without any likely interference from the 5’ CpG part of the oligonucleotide.^28,29^ We first modeled the ternary complex of CpG-STAT3D^PROTAC^ with STAT3 dimer and cereblon. Next, we included in the modeled ternary complex structure other components of the E3 ligase complex, namely DNA damage binding protein 1 (DDB1), Cullin-4A, and RING box protein 1 (RBX1) (Figure 1C, see the Methods section). Our MD simulations performed on the larger complex provided support for a potential interaction of RBX1 catalytic domain with lysine residues within the SH2 domain in the C-terminus of STAT3 protein as discussed later.

**Figure 1.**
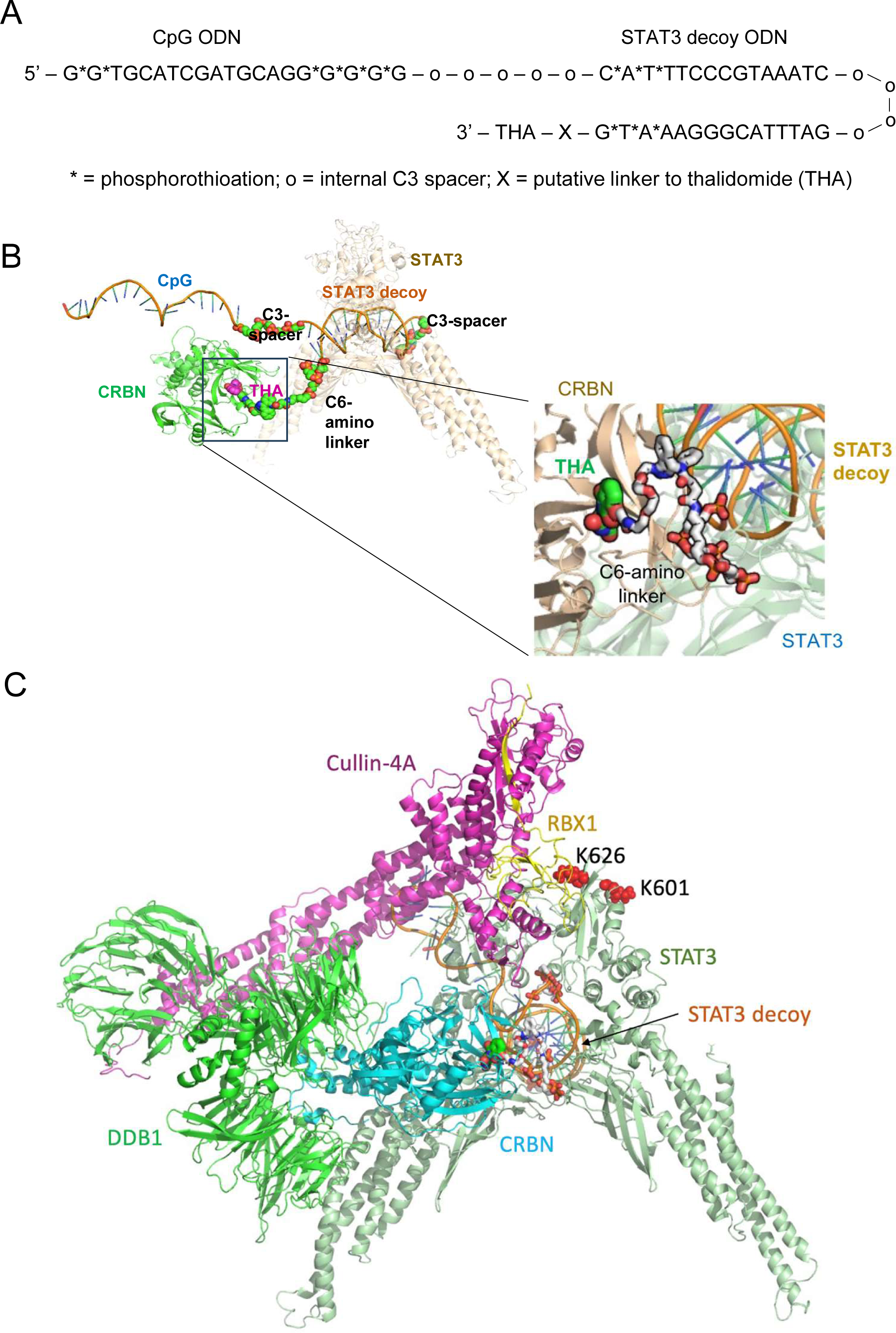
Molecular modeling of the ternary complex involving STAT3, the Oligo-PROTAC, and the E3 ligase complex in solution. **(A)** The tentative sequence of Oligo-PROTAC conjugate (C-STAT3D^PROTAC^) combining a hairpin CpG-STAT3 decoy molecule with thalidomide as an E3 ligase targeting moiety. **(B-C)** Computational molecular dynamics simulations of C-STAT3D^PROTAC^ oligonucleotide interaction with an activated STAT3 protein dimer and the E3 ligase complex displayed using Maestro software; **(B)** modeling of the thalidomide and cereblon (CRBN) binding site; **(C)** Model of the ternary complex of C-STAT3D^PROTAC^ together with the bound STAT3 proteins and the multicomponent E3 ligase complex.

Based on these modeling results, we first synthesized a thalidomide-conjugated STAT3 decoy ODN alone (STAT3D^PROTAC^) without targeting CpG domain (Figure 2A). To assess the relationship between linker length and the STAT3D^PROTAC^ activity, we compared three conjugate designs either directly conjugated to thalidomide (without linker) or connected via a single or three spacer units. The activity of STAT3D^PROTAC^ variants was assessed in mouse DC2.4 dendritic cells with constitutively activated STAT3 (DC2.4-S3C).^30^ Consistently with our computational model, the single spacer linker resulted in the maximal reduction of STAT3 protein levels in target mouse DC2.4-S3C dendritic cells (Figure 2B). Next, we performed competition experiments to verify that the inhibitory effect of STAT3D^PROTAC^ relied on both STAT3 decoy (for POI targeting) and thalidomide (for E3 ligase recruitment) parts of the conjugate. All samples were transfected with an equimolar concentration of STAT3D^PROTAC^ followed by increasing concentrations of either the unconjugated STAT3D (Figure 2C) or free thalidomide (Figure 2D), while a scrambled ODN (SCR) served as a negative control. As expected, concurrent treatments with the unconjugated STAT3D or free thalidomide almost completely abrogated inhibition at the target pSTAT3 and total protein levels by STAT3D^PROTAC^. To initially demonstrate a proof-of-concept, we transfected DC2.4-S3C (Figure 2E) and mouse A20 lymphoma cells (Figure 2F) with increasing equimolar concentrations of either STAT3D or STAT3D^PROTAC^. One day later, STAT3D^PROTAC^ reduced activated and total STAT3 levels by over 80% in DC2.4-S3C cells (at 200 nM) and A20 cells (at 400 nM) (Figure 2EF). Finally, we evaluated the specificity of STAT3D^PROTAC^ inhibitory effect. As shown in Figure 2G, STAT3D^PROTAC^ effectively reduced STAT3 protein levels without affecting the closely related STAT1 and STAT5 transcription factors in the target DC2.4-S3C cells. Thus, STAT3D^PROTAC^ oligonucleotide is shown to inhibit STAT3 signaling with high selectivity.

**Figure 2.**
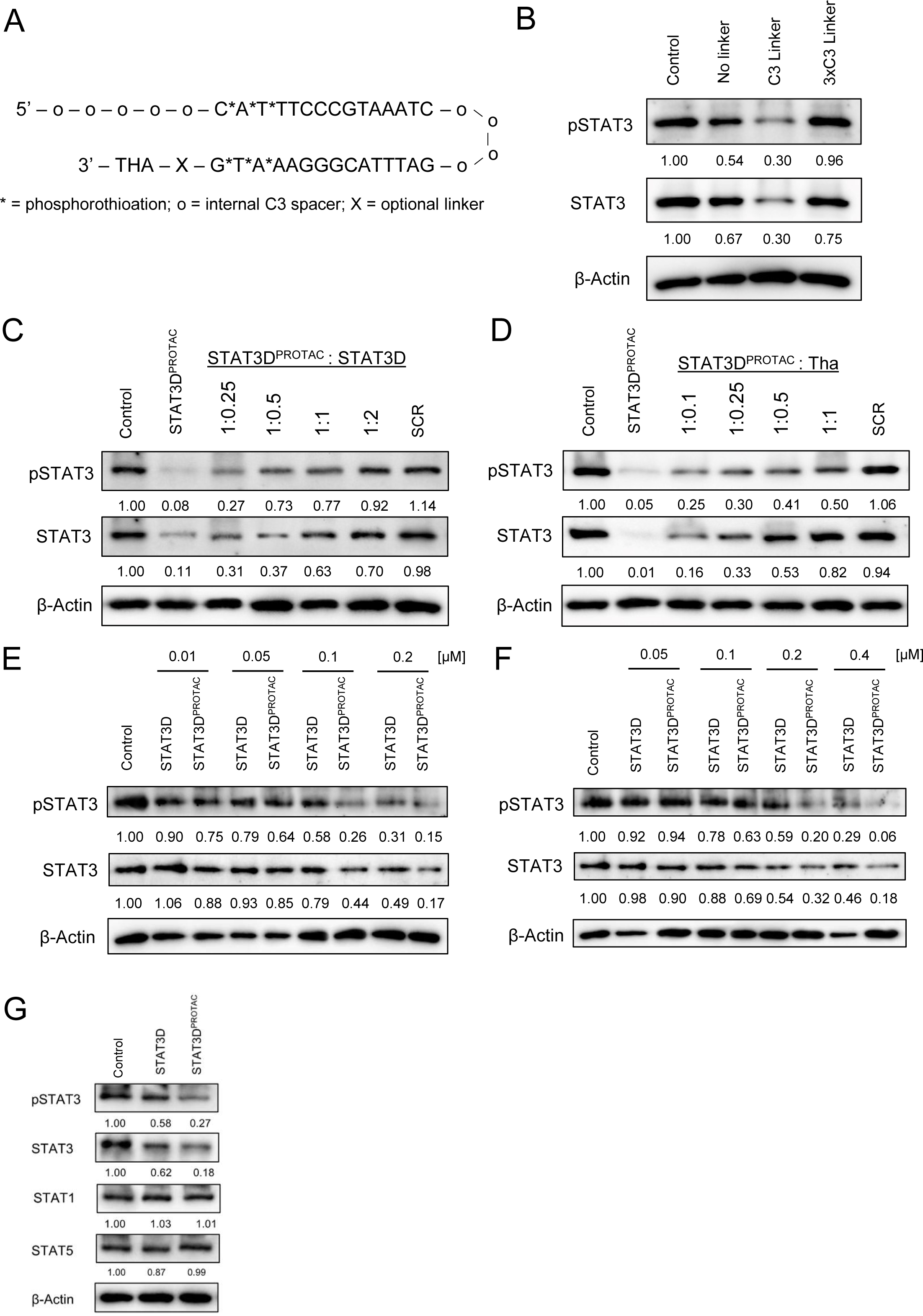
The optimization of STAT3dODN^PROTAC^ design to target STAT3 in mouse target cells. **(A)** The sequence of thalidomide-conjugated STAT3 decoy conjugate (STAT3D^PROTAC^). **(B)** Selection of the optimal linker length for tethering thalidomide to STAT3D ODN. Mouse DC2.4-S3C cells were transfected using 100 nM of the three STAT3D^PROTAC^ variants with different linker lengths. Cells were treated with IL-6 to activate STAT3 and then protein lysates were analyzed using Western blotting with β-Actin as a loading control. Levels of activated and total STAT3 were digitally quantified, normalized to β-Actin and shown as a ratio relative to the untreated sample. **(C-D)** The competition studies to assess contributions of decoy ODN **(C)** and thalidomide **(D)** moieties to the overall STAT3D^PROTAC^ activity. DC2.4-S3C cells were transfected using 100 nM of STAT3D^PROTAC^ together with increasing molar ratios of unconjugated STAT3 decoy (STAT3D) or thalidomide molecules; the scrambled ODN was used as a negative control. Levels of phosphorylated and total STAT3 protein were quantified as described and normalized to the untreated sample. **(E, F)** STAT3D^PROTAC^ reduces STAT3 activity and expression in target mouse myeloid DC2.4-S3C cells **(E)** and A20 B lymphoma cells **(F)**. Cultured cells were transfected using various concentrations of STAT3D^PROTAC^ or STAT3D alone. Total and phosphorylated STAT3 protein levels were normalized to the untreated sample and quantified as before. **(G)** STAT3D^PROTAC^ inhibits selectively STAT3 but not closely related STAT1 or STAT5. DC2.4-S3C cells were transfected using 100 nM of STAT3D^PROTAC^ or STAT3D before the evaluation of STAT protein levels. Shown are representative results from one of three repeated experiments.

### STAT3D^PROTAC^ induces CRBN-mediated proteasomal degradation of STAT3

We next verified whether STAT3D^PROTAC^ induced STAT3 inhibition is in fact dependent of proteolytic degradation of the target protein rather than its sequestration as in case of the original STAT3D molecule. As shown in Figure 3A, blocking 26S proteosome function using MG132 peptide inhibitor completely abrogated STAT3D^PROTAC^ effect and stabilized levels of activated and total STAT3 in DC2.4-S3C cells. To elucidate the mechanism further, we assessed the contribution of CRBN towards proteolytic STAT3 degradation. We compared the effect of STAT3D^PROTAC^ in CRBN-positive (Figure 3BC) and in CRBN-deficient (Figure 3BD) DC2.4-S3C cells. As expected, the CRBN-negative target cells completely lost sensitivity to STAT3D^PROTAC^ compared to CRBN-positive cells. This result confirms that CRBN recruitment by thalidomide equipped decoy ODN is critical for the inhibitory effect of STAT3D^PROTAC^. Our molecular modeling of the ternary complex involving STAT3, STAT3D^PROTAC^, and the full E3 ligase complex suggested two potential ubiquitination sites in STAT3 at lysine residues 601 and 626 which were localized in the vicinity of E2 ubiquitin binding to RBX1 (Figure 3E). Others have also suggested K601 and K626 among putative candidate ubiquitination sites within STAT3.^31^ Thus, we engineered point-mutated STAT3 protein variants with one or both lysine residues mutated to alanine and expressed in DC2.4 cells with wild-type STAT3 eliminated using CRISPR. DC2.4-STAT3KO cells were lentivirally transduced with point-mutated K601A and/or K626A variants of STAT3 and after selection, transfected using STAT3D^PROTAC^. As shown in Figure 3F, each of the ubiquitination sites seemed necessary for the proteolytic degradation of STAT3 as indicated by the loss of 80-90% of STAT3D^PROTAC^ effect in cells expressing STAT3 K601A or K626A variants (Figure 3F). The double-mutant STAT3 K601A/K626A showed a complete resistance to STAT3D^PROTAC^-induced degradation (Figure 3F). These results suggest that both lysine residues serve as non-redundant ubiquitination sites for STAT3D^PROTAC^-induced and CRBN-mediated proteolytic degradation of STAT3.

**Figure 3.**
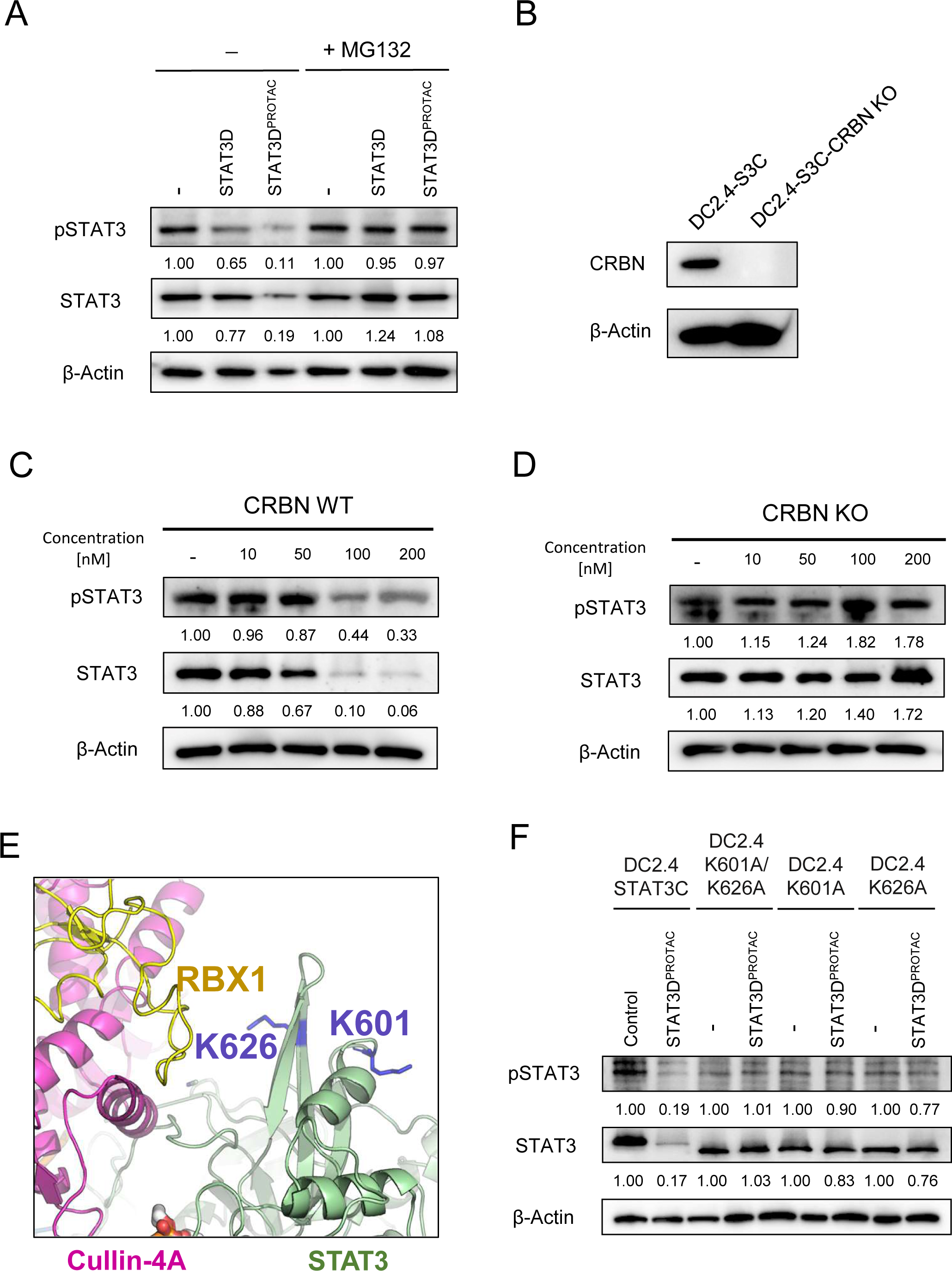
STAT3D^PROTAC^ induces CRBN-mediated proteasomal degradation of STAT3 targeting specific lysine residues. **(A)** STAT3 inhibition by STAT3D^PROTAC^ depends on proteosome activity. DC2.4-S3C cells were pretreated using 1 μM MG-132, then transfected with 250 nM of STAT3D^PROTAC^ or STAT3D alone and stimulated with IL-6 before harvesting. Total and phosphorylated STAT3 protein levels were normalized to the untreated sample and quantified. **(B-D)** STAT3 degradation by STAT3D^PROTAC^ is CRBN-dependent. DC2.4-S3C cells expressing **(B, C)** or lacking CRBN **(B, D)** were treated transfected with increasing concentrations of STAT3D^PROTAC^ or STAT3D and analyzed for STAT3 activation/protein levels using β-Actin as an internal control. **(E)** Computational modeling of the interaction between STAT3D^PROTAC^-bound E3 ligase complex, specifically RBX1 catalytic site, and STAT3 SH2 domain indicating putative ubiquitination sites at lysine residues. **(F)** DC2.4 cells stably expressing point-mutated STAT3 variants with lysine to alanine mutations at K601 and/or K626 were transfected with 100 nM of STAT3D^PROTAC^ or STAT3D alone. Cell lysates were analyzed using Western blotting and β-Actin was used as the internal control. Total and phosphorylated STAT3 protein levels compared to the untreated sample. Shown are representative results from one of three repeated experiments.

### Targeted delivery of CpG-conjugated STAT3D^PROTAC^ inhibits growth of human B-cell lymphoma xenotransplants in mice

After verifying the proteolytic and selective mechanism of STAT3D^PROTAC^ action, we equipped the 5’ end of decoy molecule with an a CpG oligonucleotide (D19) as outlined earlier (Figure 1A). The specific CpG part of complete C-STAT3D^PROTAC^ was shown to act as a targeting moiety facilitating selective uptake by TLR9-expressing immune cells, such as myeloid cells leukemia. B cell lymphoma and by the tumor-associated myeloid cells.^16,17,27^ We first tested the effect of the new C-STAT3D^PROTAC^ conjugate without any transfection reagents on mouse TLR9-positive A20 B cell lymphoma cells. A20 lymphoma was extensively tested as a target for decoy-based strategies in our previous studies.^17,18,32^ Within 24h, C-STAT3D^PROTAC^ dose-dependently reduced total STAT3 levels, with inhibition reaching maximum of 85% at 2 μM concentration (Figure 4A). In contrast, the effect of negative control treatment using an equimolar mixture of unconjugated STAT3D ODN and thalidomide was negligible (Figure 4A). As with the original STAT3D^PROTAC^, the conjugated molecule had specifically inhibited STAT3 but not STAT1 or STAT5 (Figure 2G and Supplemental Fig. S2). Next, we assessed the ability of C-STAT3D^PROTAC^ to target oncogenic STAT3 signaling in human diffuse large B cell lymphoma (DLBCL) cells. As shown in Figure 4B, C-STAT3D^PROTAC^ dose-dependently reduced activation and protein levels of STAT3 in OCI-Ly18 lymphoma cells to greater extent than the control treatment using equimolar amounts of unconjugated C-STAT3D and thalidomide, 67% vs. 24% % at 1 μM dosing, respectively. The detectable inhibitory effect of the high concentrations of reference C-STAT3D plus thalidomide treatment was likely an effect of decoy molecule interfering with autoregulation of STAT3 expression in human diffuse large B cell lymphoma cells as reported earlier.^17^ Treatment with C-STAT3D^PROTAC^ augmented downregulation of STAT3 target genes critical for lymphoma cell survival and proliferation such as *BCL2L1, CCND2, MYC* as well as proinflammatory *IL12B* compared to C-STAT3D/thalidomide (Figure 4C). To verify the potential superiority of C-STAT3D^PROTAC^ over the standard decoy design for targeting STAT3 survival signaling, we compared these two approaches in immunodeficient NSG mice bearing rapidly progressing human DLBCL. In fact, the repeated intratumoral injections of C-STAT3D^PROTAC^ (5 mg/kg) reduced tumor volume more than twice as effectively as the standard C-STAT3D (Figure 5A). Furthermore, tumors regressed in mice treated using C-STAT3D^PROTAC^ but both C-STAT3D and the negative control C-SCR injections only delayed lymphoma progression (Figure 5B). The protein analysis of whole tumors, indicated stronger inhibition of STAT3 activity by C-STAT3D^PROTAC^ than C-STAT3D, although the overall levels if STAT3 were reduced in both cases (Figure 5C). Overall, our results suggest that C-STAT3D^PROTAC^design provides superior, direct efficacy against human DCBCL over the reversible decoy inhibitor at least in the immunodeficient mice.

**Figure 4.**
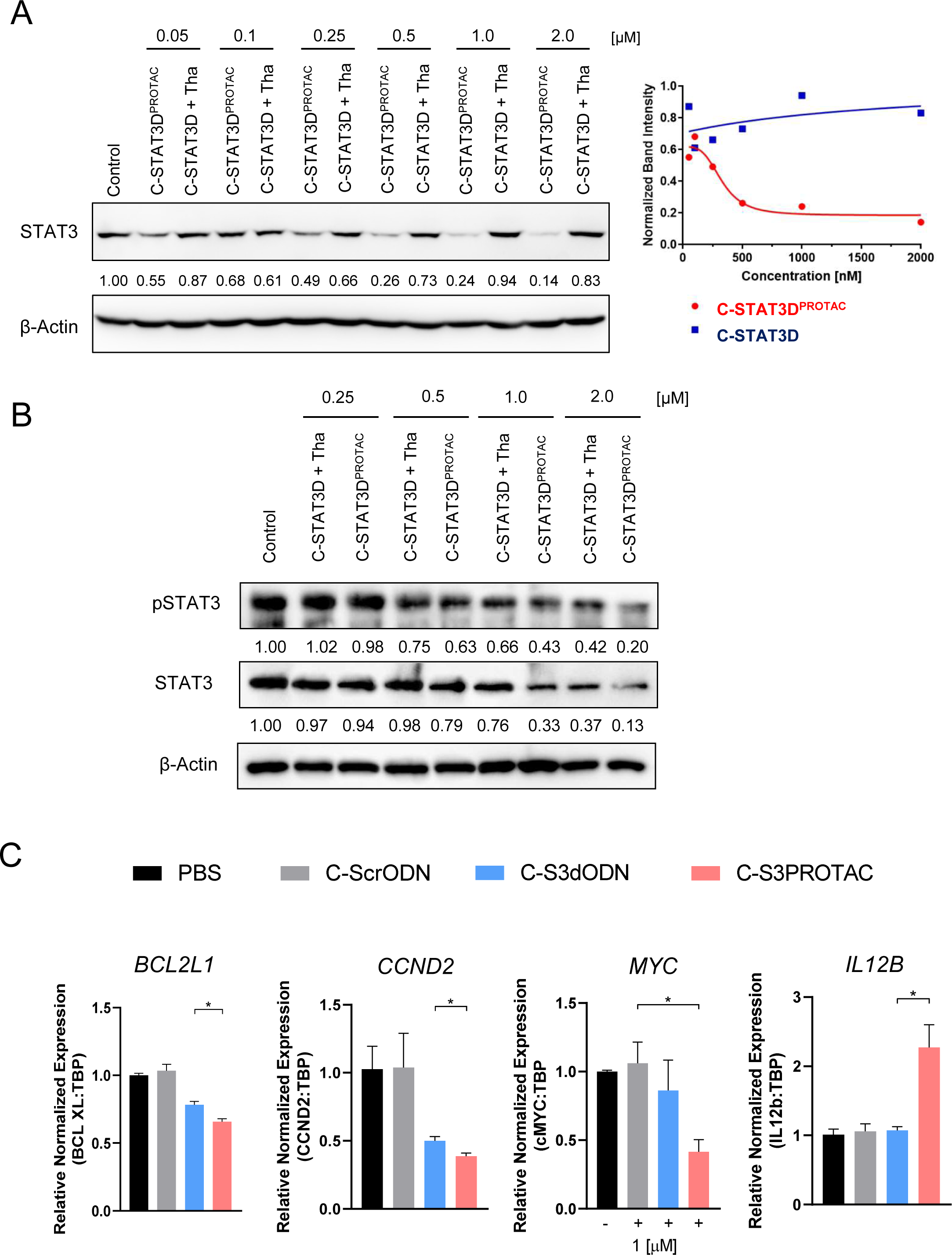
Targeted delivery of CpG-conjugated STAT3D^PROTAC^ inhibits growth of human B-cell lymphoma in vitro and in immunodeficient mice. **(A)** Naked CpG-conjugated C-STAT3D^PROTAC^ dose-dependently reduces STAT3 protein levels in target A20 B-cell lymphoma cells. Lymphoma cells were treated using increasing concentrations of C-STAT3D^PROTAC^ or an equimolar mixture of C-STAT3D with thalidomide. Total STAT3 protein levels were normalized to the untreated sample; left panel – representative Western blot results with quantification, right panel – graph of STAT3 protein levels with non-linear fit. Shown are representative data of two independent experiments. **(B)** Human OCI-Ly18 B-cell lymphoma cells were treated using C-STAT3D^PROTAC^ or C-STAT3D plus thalidomide daily for 3 days. Cell lysates were analyzed using Western blotting and β-Actin was used as internal control. Total STAT3 protein levels were normalized to β-Actin and compared to the untreated sample. **(C)** Human OCI-Ly3 cells were treated using C-STAT3D^PROTAC^, C-STAT3D plus thalidomide, or C-ScrODN over 3 days and stimulated with IL-6 before harvesting. Gene expression was examined using qRT-PCR and *TBP* as a housekeeping gene. Gene expression levels were normalized to the untreated control of OCI-Ly3 cells. Shown are means±SEM (*n*=3).

**Figure 5.**
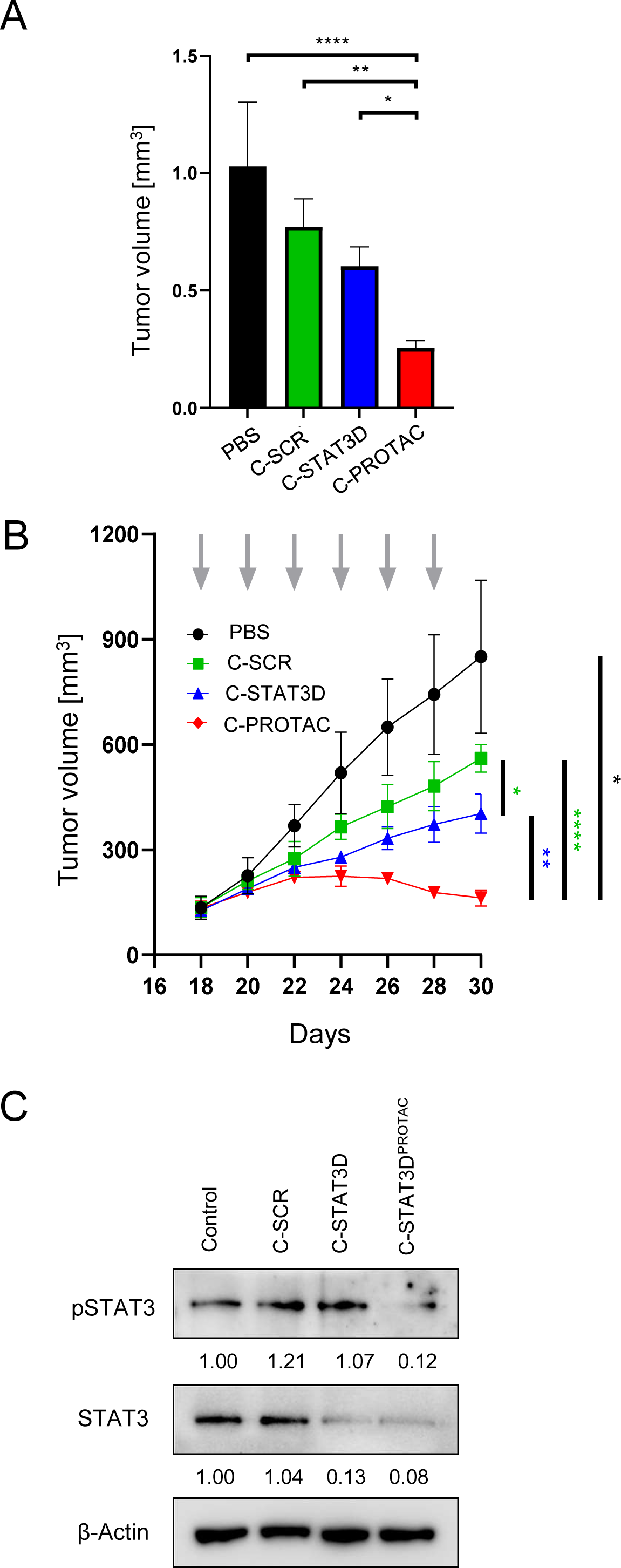
Targeted delivery of CpG-conjugated STAT3D^PROTAC^ inhibits growth of human B-cell lymphoma in immunodeficient mice. **(A-C)** Intratumoral administration of C-STAT3D^PROTAC^ inhibits growth of OCI-Ly3 B-cell lymphoma xenotransplants in immunodeficient NSG mice. 10^7^ OCI-Ly3 cells were engrafted subcutaneously and mice with established lymphomas (∼100mm^3^) were injected 6 times every other day using 5 mg/kg of C-STAT3D^PROTAC^, C-STAT3D, C-SCR ODN, or PBS. Tumors were harvested two days after the final injection to assess tumor volume **(B),** compared to tumor growth kinetics **(C)** and levels of STAT3 proteins in the whole tumors; shown are means±SEM (*n*=4).

## DISCUSSION

By combining rational design with structure-based computational analysis, we demonstrated that incorporating PROTAC activity into a cell-selective STAT3 decoy-based inhibitor can dramatically improve target inhibition and thereby the direct *in vivo* antitumor efficacy. We have previously shown that CpG-conjugated STAT3 decoy strategy results in potent immune-mediated antitumor responses against models of acute myeloid leukemia (AML) and B-cell lymphoma in immunocompetent mice.^16,17^ However, the reversibility of STAT3 inhibition by these strategies limited the direct cytotoxicity to leukemia and lymphoma cells which is an important therapeutic effect in patients’ with advanced and rapidly progressing tumors. Our results underscore the potential of using the existing decoy ODN-based oligonucleotides for the design of proteolytic degraders of undruggable TFs, such as STAT3.^2,5^ The Oligo-PROTAC design offers high molecular target specificity based on the TF-specific DNA sequence recognition, which is not affected by 3’ end modifications. As shown by the results of molecular modeling, the simplicity of Oligo-PROTAC design allows for fine-tuning the interaction between the protein target and the E3 ubiquitin ligase complex to maximize POI degradation. Furthermore, the support of modeling tools facilitates mode-of-action studies and permits identification of targeted lysine residues in the POI.

Though PROTAC technology first appeared in the early 2000s, the unique potential of targeting previously undruggable proteins through lower affinity binding has brought a burgeoning number of studies and led to recent clinical trials.^33^ Our findings represent the first demonstration of feasibility to inhibit STAT3 signaling using Oligo-PROTAC design *in vitro* and *in vivo*. While STAT3-specific PROTACs have recently been described,^26,34^ these small molecule conjugates do not offer cell-selectivity which is a crucial consideration due to the role of STAT3 in non-malignant cells, including T cells.^6,14^

Others have recently shown the potential to target oncogenic TFs, such as LEF1 and ERG, using specific double-stranded ODNs, resulting in proteolytic degradation of target proteins and reduced prostate cancer cell survival and proliferation ^35,36^. Beyond TFs, it seems feasible to employ short RNA oligonucleotides for generating PROTAC molecules targeting RNA-binding oncoproteins ^37^. Despite these rapid advances, lack of targeted delivery methods hampers further translation of these approaches to clinical application.^15,38^ While significant progress in LNP formulations of RNA enabled rapid advancement in locally administered mRNA vaccines against viral diseases and cancer, the systemic administration of oligonucleotides to organs other than liver or to specific cellular targets remains a challenge.^39^ Our study provides evidence that it is feasible to equip Oligo-PROTACs with specific targeting domains without interfering with the POI/E3 ligase complex and thereby facilitating cell-selective delivery of the naked and unformulated ODN conjugates. C-STAT3D^PROTAC^ and the original C-STAT3D utilize CpG ODN to target scavenger receptors on the variety of immune cells. These include B lymphocytes and myeloid cells, such as dendritic cells and macrophages, as well as cancer cells, e.g. AML, B-cell lymphoma cells or certain solid tumor cells in prostate cancers and glioma.^15–17^ Importantly, the interaction with endosomal TLR9 facilitates the rapid release of CpG-conjugates into the cytosol and augments potency of these oligonucleotides.^16,40^

Overall, our proof-of-concept studies on the cell-selective C-STAT3D^PROTAC^ design highlight the potential of using Oligo-PROTAC design for cancer therapy. Further optimization of C-STAT3D^PROTAC^ will focus on the key issues of molecule stability/bioavailability, immune activity and tolerability. Given that siRNA-or ASO-based CpG-STAT3 inhibitors as well as various small molecule PROTACs have reached or are near clinical testing, we believe that C-STAT3D^PROTAC^ has potential to provide safe and effective treatment for patients with B-cell lymphoma, and potentially other cancer indications.

## MATERIALS AND METHODS

### Cells

Human OCI-Ly3 B cell lymphoma line was purchased from Deutsche Sammlung von Mikroorganismen und Zellkulturen (DSMZ). Human OCI-Ly18 cells were provided by Dr. Larry Kwak (City of Hope/COH, CA, USA). Mouse dendritic DC2.4 cells were originally from Dr. Kenneth Rock (University of Massachusetts Medical School, MA, USA). Mouse A20 B cell lymphoma line was purchased from American Type Culture Collection (ATCC, Manassas, VA, USA). DC2.4, A20, OCI-Ly3 and OCI-Ly18 cells were cultured in RPMI1640 with 10-20% fetal bovine serum (FBS). To generate DC2.4 cells with constitutively activated STAT3C (DC2.4-S3C) ^30^ or point-mutated K601A and K626A STAT3 variants, expression plasmids were designed and purchased from VectorBuilder (Chicago, IL, USA), then cloned into a third-generation lentiviral vector (pMDLg/pRRE/pRSV-Rev/pMD2.G). The STAT3 mutation-bearing lentiviral vectors were then transduced into DC2.4-CRISPR-STAT3KO cells and mutant cells were selected for with puromycin and GFP^+^ cells were sorted. All cells were regularly tested for mycoplasma contamination using the LookOut mycoplasma PCR detection kit (Sigma-Aldrich, St. Louis, MO, CA).

### Mice

All animal experiments were carried out in accordance with established institutional guidance and approved protocols from the institutional animal care and use committee (COH, Duarte, CA, USA). *NOD/SCID/IL-2RγKO* (NSG) mice, originally obtained from the Jackson Laboratory (Bar Harbor, ME, USA), were maintained at COH. Mice were injected subcutaneously with 10^7^ OCI-Ly3 cells in PBS and lymphoma engraftment and progression were monitored by caliper measurements.

### Oligonucleotide Design

All of the following oligonucleotides were synthesized in the DNA/RNA Synthesis Core (COH) as previously described^27^ and then conjugated to modified thalidomide moiety using click chemistry as shown in the Supplemental Figure 1. The resulting conjugates are illustrated below (o = internal C3 spacer, X = 3’-C6-amino linker, * = phosphorothioation, THA = thalidomide).

#### C-STAT3dODN

5’-G*G*TGCATCGATGCAGG*G*G*G*G – o – o – o – o – o – C*A*T*TTCCCGTAAATC – o – o – o – o – GATTTACGGGAA*A*T*G-3’

#### C-scrODN

5’-G*G*TGCATCGATGCAGG*G*G*G*G – o – o – o – o – o – A*C*T*CTTGCCAATTAC – o – o – o – o – GTAATTGGCAAG*A*G*T-3’

#### C-STAT3dODN^PROTAC^ (No Linker)

5’-G*G*TGCATCGATGCAGG*G*G*G*G – C*A*T*TTCCCGTAAATC – o – o – o – o – GATTTACGGGAA*A*T*G – THA-3’

#### C-STAT3dODN^PROTAC^ (Single Linker Unit)

5’-G*G*TGCATCGATGCAGG*G*G*G*G – o – o – o – o – o – C*A*T*TTCCCGTAAATC – o – o – o – o – GATTTACGGGAA*A*T*G – o – THA-3’

#### C-STAT3dODN^PROTAC^ (Three Linker Units)

5’-G*G*TGCATCGATGCAGG*G*G*G*G – o – o – o – o – o – C*A*T*TTCCCGTAAATC – o – o – o – o – GATTTACGGGAA*A*T*G – o – o – o – THA-3’

### Modeling of the STAT3-PROTAC-CRBN complex

The crystal structure of oligo-decoy bound STAT3 was downloaded from the PDB databank (ID: 1BG1).^41^ The structure was prepared using the Protein Preparation Wizard^42^ in Maestro by adding hydrogen atoms and missing residues, followed by minimization using MacroModel^43^. The CpG and linkers were attached to the oligo-decoy using molecular modeling in Maestro. Lenalidomide bound CRBN structure was downloaded from the PDB databank (ID: 5FQD)^44^ and placed close to the PROTAC linker of STAT3 using rigid body transformations in Maestro. Thalidomide was modeled by modifying the bound lenalidomide, followed by the addition of covalent bond with the C6-amino linker. The resulting CRBN/PROTAC bound STAT3 structure was parameterized in AMBER16^45,46^ using the FF14SBonlySC^47^ and Parmbsc1^48^ forcefields for protein and nucleic acid segments respectively. The linkers and thalidomide moieties were parameterized using the GAFF2 forcefield.^49^ Partial charges were obtained by fitting a restrained coulomb function to the electrostatic potential (RESP)^50^ obtained using JAGUAR^51^. The partial charges were calculated using the online R.E.D, server.^52^ The system was optimized by equilibrating for 100 ns at 290K using the IGB8 Generalized Born implicit solvation model^53^ and AMBER MD simulation program.

### Modeling of the STAT3/PROTAC bound E3 ubiquitin ligase

The crystal structure of CRBN bound to DDB1, Cullin4A and RBX1 was downloaded from the PDB databank (ID: 2HYE).^54^ This structure was then aligned to the CRBN bound STAT3 complex using PyMOL.^55^ Finally, the DDB1, Cullin4A and RBX1 moieties from the first structure were combined with the CRBN bound STAT3 to obtain the entire E3 ligase complex.

### Transcriptomic and Protein Assays

For quantitative PCR (qPCR), total RNA was extracted from cultured cells using the Maxwell RSC simplyRNA Cells system (AS1390, Promega, Madison, WI, USA), then reverse transcribed into cDNAs with the iScript cDNA synthesis kit (Bio-Rad). The qPCR was then carried out using specific primers for *BCL2L1*, *CCND2*, *MYC*, *IL12B*, and *TBP* as previously described ^56,57^ with a CFX96 Real-Time PCR Detection System (Bio-Rad). Western blots were performed and described previously ^56^ using antibodies specific to pSTAT3, STAT3, STAT1, STAT5 (Cell-Signaling Technology, Danvers, MA, USA) or β-Actin-HRP (Sigma-Aldrich, St. Louis, MO, USA). Blots were imaged in a Bio-Rad ChemiDoc MP System using enhanced chemoluminescence (ECL; SuperSignal West Femto Maximum Sensitivity Substrate), and the resulting images were analyzed with accompanying Bio-Rad Image Lab software and GraphPad Prism 8. Cytokine levels in cell culture supernatants were measured using the Luminex system.

### Statistical Analysis

An unpaired t-test was used to determine the statistical significance of differences between two treatment groups. Two-way analysis of variance (ANOVA) plus Bonferroni post-tests were utilized to estimate the statistical significance of differences between multiple treatment groups. The relationship between two groups was determined via correlation and linear regression. The p values are indicated in figures with asterisks: *p < 0.05; **p < 0.01; ***p < 0.001; ****p < 0.0001. Data were analyzed with Prism v.8.4.3 software (GraphPad).

## DATA AVAILABILITY STATEMENT

All data and reagents generated within this study are available from the corresponding author upon a reasonable request.

## ACKNOWLEDGEMENTS

We are grateful to the staff at Analytical Cytometry, DNA/RNA Synthesis, Analytical Pharmacology and Animal Resource Cores (COH). This work was supported in part by the National Cancer Institute/National Institutes of Health awards number R01CA213131, R01CA284593 (M.K.), P50CA107399 (S.F.) and P30CA033572 (COH). The content is solely the responsibility of the authors and does not necessarily represent the official views of the National Institutes of Health.

## AUTHORS CONTRIBUTIONS

Conceptualization, M.K., P.S. and N.V.; Methodology, Z.Z.,J.H., S.B., D.W.; Investigation, J.H., Z.Z., Y.L. and D.W.; Writing – Original Draft, Z.Z., J.H. and M.K.; Writing – Review & Editing, M.K.; Funding Acquisition, M.K., L.K., S.F.; Resources, P.S., M.K.; Supervision, M.K.

