## Supplemental Data for "Oligo-PROTAC strategy for cell-selective and targeted degradation of activated STAT3"

Supplemental Figures

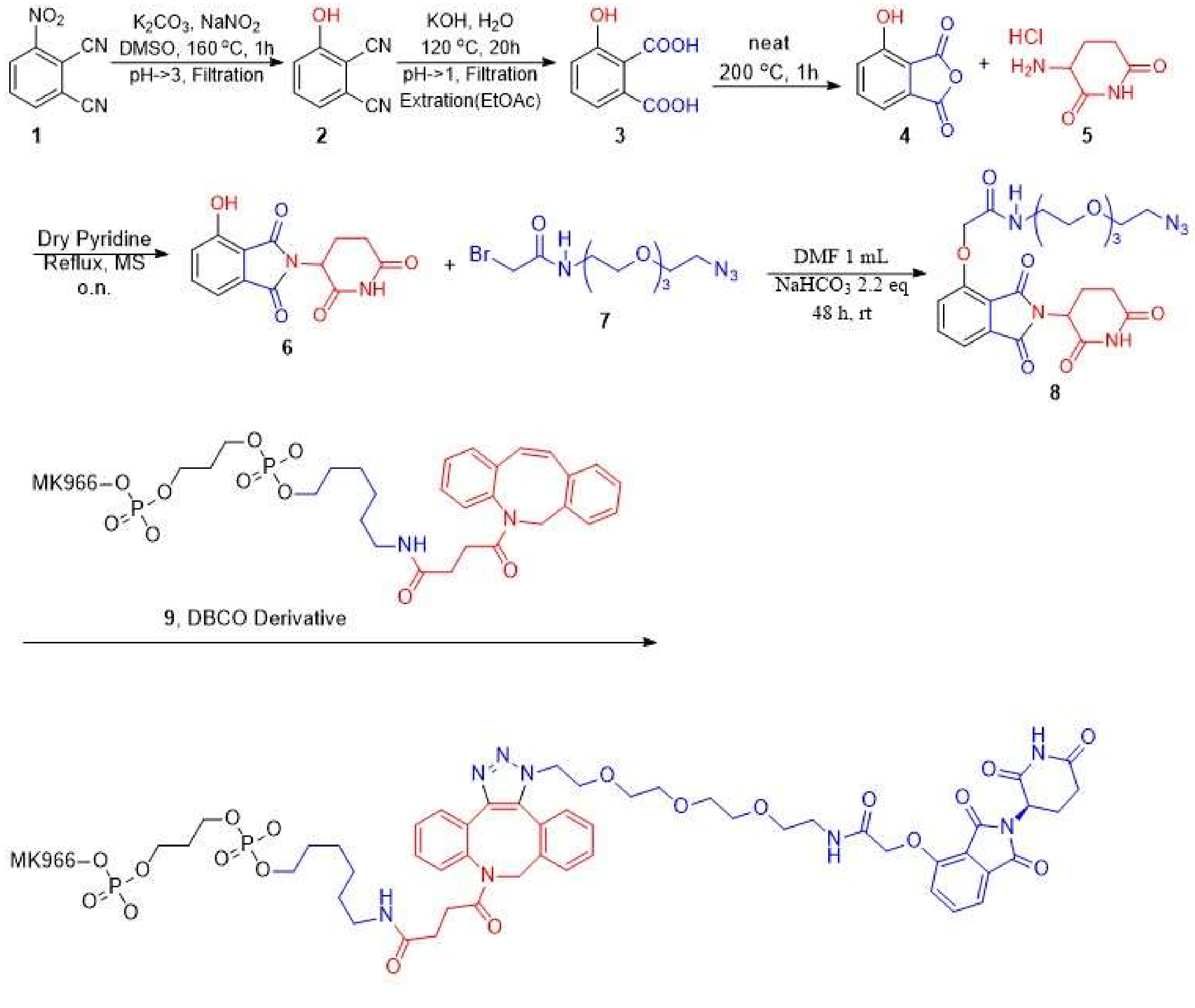


10, MK966 Thalidomide Conjugates

**Supplemental Figure S1.** Schematic representation of major steps in the chemical synthesis of C-STAT3D$^{PROTAC}$ conjugate.

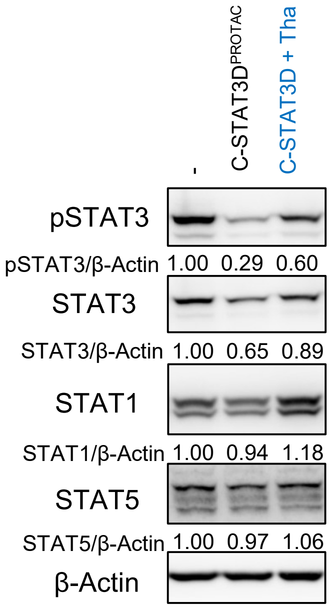


**Supplemental Figure S2.** C-STAT3D$^{PROTAC}$ induces selective degradation of STAT3. DC2.4 cells were treated using 500 nM C-STAT3D$^{PROTAC}$ or C-STAT3D plus thalidomide daily for 48h and stimulated using 20 ng/mL mouse IL-6 6 h before harvesting the cells. pSTAT3, STAT3, STAT1 and STAT5 levels were assessed using Western blotting, quantified and normalized to actin and compared to the IL-6 treated control.

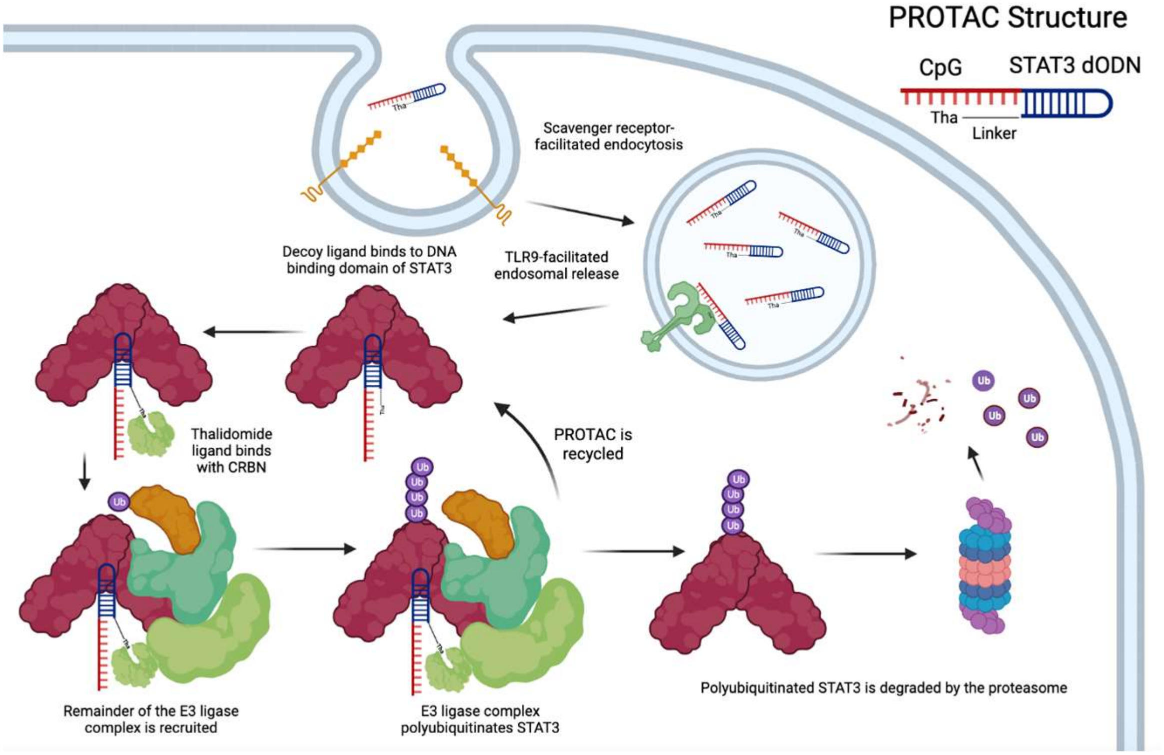


**Graphical Abstract.** Immune cell-selective C-STAT3D$^{PROTAC}$ induces proteolytic degradation of STAT3; Ub – ubiquitin, The – thalidomide moiety; CpG – cytosine-phosphate-guanosine motif; dODN – decoy oligodeoxynucleotide; TLR9 – Toll-like receptor 9.
